# Many morphs: parsing gesture signals from the noise

**DOI:** 10.1101/2023.08.07.551967

**Authors:** Alexander Mielke, Gal Badihi, Kirsty E. Graham, Charlotte Grund, Chie Hashimoto, Alex K. Piel, Alexandra Safryghin, Katie E. Slocombe, Fiona Stewart, Claudia Wilke, Klaus Zuberbühler, Catherine Hobaiter

## Abstract

Parsing signals from noise is a general problem for signallers and recipients, as well as for researchers studying communicative systems. Substantial research efforts have been invested in comparing how other species encode information and meaning in their signals, and how signalling is structured. However, our ability to do so depends on identifying and discriminating signals that represent meaningful units of analysis. Early approaches to defining signal repertoires applied top- down approaches, classifying cases into predefined signal types. Recently, more labour-intensive methods have taken a bottom-up approach describing the features of each signal in detail and clustering cases into types based on patterns of similarity between them in multi-dimensional feature-space that were previously undetectable. Nevertheless, it remains essential to assess whether the resulting repertoires are composed of relevant units from the perspective of the species using them, and redefining repertoires when additional data makes more detailed analyses feasible. In this paper we provide a framework that takes data from the largest set of wild chimpanzee (*Pan troglodytes*) gestures currently available, splitting gesture types at a fine scale based on modifying features of gesture expression and then determining whether this splitting process increases the information content of the communication system. Our method allows different features of interest to be incorporated into the splitting process, providing substantial future flexibility across - for example - species, populations, and levels of signal granularity. In doing so we provide a powerful tool allowing researchers interested in gestural communication to establish repertoires of relevant units for subsequent analyses within and between systems of communication.

## Introduction

If you want to see a biologist struggle, ask them to define what a species is. None of the easy answers apply to the real world. We spend a substantial amount of our time trying to fit messy, highly variable data into tidy, artificial boxes. In practice, we know that perfectionism is largely futile and accept that, at best, we are describing a fraction of the variation we see expressed in the world around us. Nevertheless, our work is often founded on comparison - we fit data into groups to be able to interrogate them. We classify individuals into species, populations, sexes, age groups. We classify behaviour using carefully constructed ethograms (Bateson & Martin, 2021). In practice the expression of all of these is graded (Hey et al., 2003), and we can choose different levels of detail – lumping or splitting sets – depending on our question of interest (or on the data available).

A clear example of this process is the study of communication in non-human species. First, we assign signals to distinct channels such as vocalisations and gestures, despite use of these channels overlapping extensively (Slocombe et al., 2011). We then take the rich, graded systems expressed across those channels and carve them up into sets or ‘repertoires’ based on our intuition and ability to perceive and categorise streams of graded or blended signals into discrete units (Parr et al., 2005; Hobaiter & Byrne, 2017). This parsing process is something the human brain is particularly good at (Saffran et al., 1996) – spoken human language as well as language-accompanying gestures and facial signals are produced not as a set of discrete signals, but as a continuous stream of information that is perceived discreetly (Jack et al., 2014). Doing so during early language learning is automatic: we are rarely aware that we are rapidly processing an extremely rich set of statistical cues (Saffran et al., 1999; Swingley, 2005).

When we rely on this intuitive classification to ask questions about animal communication – about information and meaning, structure and syntax, individual variation and community or species-wide shared characteristics – our results become conditional on the choices we make in categorising signals, opening the door for considerable biases due to researcher degrees of freedom (Wicherts et al., 2016). Historically, in non-human primate (hereafter primate) vocal and facial communication, both highly graded systems, we often started by deriving minimal sets of stereotypical signals (e.g., ’hoo’) that were then further qualified by context (‘resting hoo’, ‘alarm hoo’, ‘travelling hoo’; e.g., Marler, 1976). Doing so typically involved discarding significant portions of data where signals fell outside of classification boundaries - for example, chimpanzee (*Pan troglodytes*) ‘neutral’ facial signals contain a lot of subtle muscle activation (Parr & Waller, 2006). As a result, we may have underestimated the actual amount of information being transmitted. For example, the ‘Silent Bared Teeth’ display in crested macaques (*Macaca nigra*) was recently shown to comprise at least four distinct forms that varied reliably with context (Clark et al., 2020). Similarly to ‘Silent Bared Teeth’ faces, early descriptions of unified call types have been slowly split into more detailed repertoires based on acoustic features and differences in use (e.g., Clay et al., 2015; Crockford et al., 2018; Slocombe & Zuberbuhler, 2005; Slocombe et al., 2007). Advances in detailed acoustic analysis introduced a more data driven and replicable approach to studies of vocal communication (Sainburg et al., 2020). Critically, only in some cases subsequent playback experiments have confirmed that listeners meaningfully distinguish between these call variants (Slocombe et al., 2009; Slocombe et al., 2010). For many species, particularly those with highly graded call systems, the exact number of call types in the repertoire has remained elusive, with variation increasingly described using fuzzy element boundaries and clustering algorithms (Fischer et al., 2017; Wadewitz et al., 2015).

Researchers traditionally approached primate gestures differently, in that the system was historically split into many more distinct units than the vocal or facial channels, with little additional detail about signal production, so signals appeared less graded and more like a rich repertoire of easily differentiated units or ‘gesture types’ (Plooij, 1979; Tomasello et al., 1985). The difference between discrete gesture types has usually been established based on a mixture of morphological and contextual definitions designed to achieve high reliability within studies, but less consistency between studies (Rodrigues et al., 2021). This process also potentially splits gesture units along arbitrary lines where features salient to a human coder may not be detected or relevant to the individuals using them. For example, partner contact with fingers (‘Poke’), palm (‘Slap’), or fist (*‘*Punch*’*), are associated with overlapping patterns of goals in chimpanzees (Hobaiter & Byrne, 2017), despite sometimes being coded as discrete ‘types’ by human observers (Hobaiter & Byrne, 2011). Conversely, leaf-clipping is often lumped by coders, but when we split them according to features such as the location of the tearing action on the leaf, different variants become apparent (Badihi et al., 2023). More recently, there has been a push for replicable definitions of gesture types, and a focus on bottom-up detailed coding to allow for more granular analyses of gestural communication. For example, the use of ‘Minimum Action Units’ or MAU (Grund et al., 2023), distinguishes the initial physical movement of the gesture from the hold or repetition phases. Original coding is highly split with systematic detailed descriptions of gesture form, i.e., MAU, alongside gesture modifiers (Grund et al., 2023). We call these additional morphological characteristics of a gesture action ‘modifiers’ (termed ‘features’ in previous publications, e.g., Hobaiter & Byrne, 2017), as they modify the appearance of the gesture action. While labour intensive, doing so allows researchers to define gesture units flexibly, and at different levels of granularity depending on features of interest for a particular analysis or on the characteristics of available data. While potentially powerful, there remains one substantial challenge to effectively applying this method to the analysis of gestural data: discriminating levels of lumping or splitting of gesture tokens that provide salient differences in information to the primates using them.

One hurdle in exploring gradation in gestural repertoires is that the levels of lumping and splitting vary not only between different repertoires, but also between units within them. Each signal is typically described as a set of physical characteristics – the defining movements of the gesture action plus a set of modifiers. But in some cases, modifiers are already used to split gesture actions. For example, in the literature, some units are made of gesture actions split based on duration (e.g., long and short touches; Fröhlich et al., 2016), body part (e.g., embraces with arms or feet; Liebal et al., 2004), repetition (e.g., hitting once vs repeatedly; Hobaiter & Byrne, 2011) or the use of one or both limbs (e.g., grabbing with one or two hands; Genty et al., 2009). At the same time, for other gesture units in the same repertoire, these same properties might be lumped and considered to be variation within the expression of a single gesture unit. Sometimes, some – but not all – units are split by their context of production (e.g., chest beat vs chest beat play, Genty et al., 2009; lunging vs lunging in play, Gupta & Sinha, 2019). In more commonly produced actions, patterns of difference that could be used to discriminate separate gesture units may be more easily detected than in rarely observed ones. But, typically, the approaches to defining a novel gesture unit via lumping and splitting are not transparent (Rodrigues et al., 2021). Importantly, there is no absolute correct level of splitting or lumping, the level of detail in a communicative repertoire depends on the question being asked. In spoken language, we construct sets of phonemes and syllables, as well as of words and higher order grammatical structures. However, it is important that a set is composed of similarly characterised units – whereas our current ape gesture repertoires are frequently composed of a mix of the equivalent of syllables and words.

Gradedness in gestures might not be as apparent to observers as it is in vocal and facial communication. However, the actions that make a gesture recognisable can be produced with different morphological features: they can be produced in combinations of different channels (acoustic, visual, tactile), with different limbs, at different intervals, at different speeds, and using different space. For example, an Embrace can be defined as the signaller wrapping their limbs around the recipient, making it distinct from any other gesture action. However, whether one or both arms are used or the feet (Liebal et al., 2004), which arm, which body part is embraced, for how long, etc., may still contain information about the goal of the gesture, the context of use, and/or the social relationship between signaller and recipient. While ‘Embraces’ are clearly distinct gesture actions that most observers can reliably identify, a more fine-grained splitting process (e.g., one-armed embraces from the side around the shoulders vs bimanual embraces from behind) is possible. A recent study identified 36 different forms of the ‘touch’ gesture in chimpanzees, directed toward 70 different locations on the body (Bard et al., 2019). Typical modifiers include the body part with which the signaller produced the gesture; the body part on the recipient where the gesture made contact; whether the gesture was repeated; which limb was used; whether specific body parts (such as the fingers, wrist, or elbow) were flexed; and whether an object or physical substrate was involved (Bard et al., 2019; Grund et al., 2023; Hobaiter & Byrne, 2017; Roberts et al., 2012). Some coders ascertain the actions of multiple body parts for each action (e.g., Forrester, 2008), but this can be time-intensive when generating large datasets (Cartmill et al., submitted), at least until automation through pose-estimation and movement-tracking programs such as DeepLabCut become possible (Wiltshire et al., 2023). It is important to use existing large datasets to establish which level of granularity and coding effort is both useful and justifiable.

As soon as we start to compare communities or species, analyse meaning, or analyse sequential structure, the ability to discriminate boundaries on the basis of relevant variation, to parse gesturing into discrete gestural units, becomes of substantial importance. A previous study attempted to use unsupervised cluster analysis to establish types of gestures based on morphological features of each gesture instance alone without resorting to human-defined gesture actions (Roberts et al., 2012). However, the large number of possible morphological features in relation to the small number of available data points made it impossible to interpret the results of early efforts. As gesture datasets are coded manually, small sample sizes often prevent meaningful statistical analyses, and splitting gesture actions with too much detail leads to large numbers of rare combinations of modifiers, amplifying this challenge (Bard et al., 2019). One study of chimpanzee gestures tried to define which features in a highly-split repertoire were salient on the basis of whether they appeared to shift the signallers’ goals (for example, whether alternate or simultaneous hitting actions led to a different outcome; Hobaiter & Byrne, 2017). A first attempt to take a species-centric perspective to defining gestural repertoires, this approach was largely descriptive. Moreover, the assessment of whether to lump up from the finest level of splitting was made based on similarity of the signaller’s goal across the different units – for example, where two Hit actions were made with different body parts, these would be lumped up if they achieved a similar goal or set of goals. In incorporating signaller goal as a feature that discriminated gesture ’type’, did not allow us to subsequently evaluate whether the resulting signal sets were more or less ’meaningful’ – in terms of goal specificity – to the chimpanzees using them (Hobaiter & Byrne, 2017).

In this paper, we address the challenge of identifying gradedness in gestural communication by designing a process that lets us split existing gesture actions at a fine scale and determine whether this splitting process increases the information content of the system. This approach builds on the bottom-up, highly detailed coding scheme employed to generate the data (Grund et al., 2023). We take a pragmatic approach - following gesture coding traditions in human co-speech and co-sign gesture (Kendon, 2004), we retain the traditional concept of a predefined set of gesture actions, as a useful shorthand summarising a large number of possible dimensions that could otherwise create a lot of noise. Similarly to the integration of handshape, location, and movement to discriminate signs in American Sign Language (Stokoe, 1960), we incorporate information from a series of modifiers. We use unsupervised clustering to split each gesture action into multiple variants based on the modifier distributions but introduce sampling thresholds to prevent the creation of a near-infinite number of rare cases. This procedure provides us with a maximum repertoire (given the available data) that has been created without including the interaction’s social context or outcome. This key information on context and outcome can then instead be used to establish which of the variants of each gesture action – which we term ‘gesture morphs’ (Grund et al., 2023) – represent distinct communicative signals that contain information for recipients.

Even with our understanding of the language space, our ability to classify words in new languages without input from speakers takes huge datasets (Hvitfeldt & Silge, 2022). Nevertheless, increasing computational power, larger datasets, and sophistication in machine learning and model-based analysis have offered new methods to approach this problem. Parsing words and sentence boundaries out of streams of speech data is a fundamental task of natural language processing – a task which supervised and unsupervised machine learning models are increasingly capable of solving efficiently (Chollet et al., 2022). For example, human facial movement recognition is similarly able to identify relatively discrete facial signals out of video data (Zhou et al., 2010). Applying similar approaches to animal calls has allowed for the data-driven generation of population- or species-specific repertoires, especially for highly vocal species with large training datasets available (e.g., orcas: Bergler et al., 2019; budgerigars and long-billed hermits: Keen et al., 2021; meerkats: Thomas et al., 2022). More recently, the availability of standardised software for feature extraction and analysis, such as Koe (Fukuzawa et al., 2020) and DeepSqueak (Coffey et al., 2019), makes these tasks easier than ever where large corpora are available. Further, for species with highly graded call systems, fuzzy clustering approaches might prove more insightful in quantifying the complexity of the call repertoire (Fischer et al., 2017). Automated analyses of repertoires of animal facial signals have been introduced (Dolensek et al., 2020) but are currently limited to reactions to specific ‘emotional’ stimuli (such as the detection of pain; Andersen et al., 2021) and are hampered by the lack of large datasets of natural facial signals.

Cluster detection of any kind has only been applied in gesture to insufficiently large datasets (e.g., n=128 tokens, Roberts et al., 2012), limiting interpretation. Cluster analysis is data-intensive and any result is conditional on amassing a) a sufficiently large dataset (Bouveyron et al., 2019), and b) one that is coded in a bottom-up transparently structured way, both of which are extremely time-intensive. Gestural video data, especially when collected in non-standardised ‘noisy’ visual-audio environments such as forests, are higher-dimensional than sound files and videos of faces (which can be treated as two-dimensional without losing much information), introducing considerable challenges for automated feature extraction. Here we take advantage of a newly available large set of gestural data from East African chimpanzees (*Pan troglodytes schweinfurthii*): the Gestural Origins database (Hobaiter et al., 2021), generated with the GesturalOrigins coding scheme (Grund et al, 2023). We use this dataset to test whether gestural actions, the most typical level of discrimination in a gestural repertoire, can be split further through the systematic application of modifiers, and whether such a split would add information to investigations of the function and form of gestural communication. We use cluster detection to determine divisions within gesture actions that take into account the dependence structure of modifiers that occur at least five times, indicating that these splits are meaningful, replicable using other datasets, and can be used for further analysis. We hope that in doing so we offer a systematic framework for the classification of gestural data. Importantly, the level of granularity in any given study will depend on available data – the more data available (overall, and for specific gesture actions), the more splitting is possible. Using the largest existing gesture dataset, we provide a suggestion for the dimensions along which to split gesture actions, and test whether this approach can improve our understanding of gesture meaning.

## Methods

### Dataset

To detect the underlying structure of gesture actions, we focus on the well-studied gestural system of East African chimpanzees (*Pan troglodytes schweinfurthii*). We combined data from five long-term research communities in four sites: Sonso and Waibira communities, in the Budongo Forest Reserve, Uganda; the Issa Valley chimpanzees, in Western Tanzania; the M-group of Kalinzu Forest chimpanzees, Uganda; and the Kanyawara community, in the Kibale National Park, Uganda. Details of the observation effort and sample size for each community can be found in Table 1. Data were coded using the same protocol across communities. In total, 7,879 gestures were available for which coding was complete and signaller and recipients could be identified, representing 90 distinct gesture actions before preprocessing. The dataset is biased towards the Budongo chimpanzees, which provided 82% of all gestures to the dataset, and within Budongo to chimpanzees in the Sonso community, which provide 62% of all gestures to the dataset. Gesture data were collected across a full range of behavioural contexts (n = 22, see Grund et al., 2023 for ethograms), but 31% of gestures occurred before or during play.

**Table 1:** Description of the communities.

| Community | Number of<br>Gesture<br>Tokens | Number of<br>Gesture<br>Tokens - Final | Number of<br>Signallers | Gesture<br>Actions |
| --- | --- | --- | --- | --- |
| Sonso | 4956 | 4903 | 75 | 48 |
| Waibira | 1537 | 1477 | 84 | 41 |
| Issa | 586 | 576 | 32 | 40 |
| Kanyawara | 494 | 493 | 40 | 30 |
| Kalinzu | 306 | 299 | 41 | 27 |

The coding scheme for this project has been described in detail in Grund et al. (2023), including definitions of all gesture actions and modifier variables employed to describe the full dataset. To study the variation in modifiers within gesture actions, we selected four of these modifiers: (i) the body part with which the gesture was produced (‘Body Part Signaller’; 11 levels); (ii) the body part with which contact was established either on the recipient or the signaller’s own body (‘Body Part Contact’; 9 levels); (iii) whether the gesture was repeated rhythmically or not (‘Repetition’; 2 levels); and (iv) whether the gesture was produced with one limb, both limbs at the same time, or both limbs alternatingly (‘Laterality’; 3 levels). More information about the different modifier levels and Interobserver reliability tests on gesture coding can be found in the Supplementary Materials. Initially, we had included the involvement of objects in a gesture as a modifier – however, the original list of gesture actions were largely pre-split based on object use (e.g., ‘Hit Object’ was already discriminated from ’Hit Other’ at the gesture action level), thus there was little variation within gesture actions. We had the choice of either lumping up the actions and retaining object involvement as a modifier, or splitting object involvement at the level of the gesture actions. Here we decided to do the latter, incorporating it into the gesture action level, as this tended to more closely map on to previous repertoires and simplified our set of modifiers. The relevance of object-use as a modifier might be different if this approach was applied to a different dataset, for example chimpanzees may vary their use of objects depending on their structural and acoustic properties (Fitzgerald et al., 2022; Gibson et al., 2023). Other modifiers are available in the original dataset but are often only applicable to a subset of gesture actions or cases within gesture actions (for example: finger flexion); or are rarely or never part of other primate gesture coding datasets, thereby limiting applicability of results for other users. We focused on those modifiers that are commonly coded across research groups and that affect most gesture actions. Some modifiers were originally coded with more levels than included here but were lumped based on predetermined criteria - for example, left- and right-handed unimanual gestures were combined as ‘unimanual’.

In the original coding protocol, some gesture actions were pre-split based on modifiers that are of interest here, resembling predefined morphs. For example, ‘Hit Other’ indicates that the recipient received one hit, while ‘Hitting Other’ indicates that multiple hits had taken place. To reduce all gesture actions to the same level of splitting before determining the morphs and checking whether these predefined splits were justified, we lumped gesture actions that were pre-split in the original coding based on modifiers of interest (here both would be lumped as gesture action ‘Hit Other’ with modifier repetition as yes (Hitting) or no (Hit)) before proceeding with the analysis, resulting in 61 gesture actions across all sites.

### Preprocessing

Rare occurrences of gesture actions or modifier levels make it hard to understand the underlying usage rules - a common problem in linguistic analyses that is usually resolved by excluding rare elements (Levshina, 2015). Thus, in an initial preprocessing step, we removed 19 of the 61 gesture actions that had fewer than 10 occurrences available in the dataset, assuming that it is currently impossible to discriminate distinct morphs within them.

In addition to having a filter for gesture actions, we included several inclusion criteria for modifiers: First, within any gesture action, modifier levels that occurred fewer than five times were combined into an ‘Other’ category (Hvitfeldt & Silge, 2022); if the ‘Other’ category had fewer than five cases, those cases were set to missing values. Second, if a modifier was coded as ‘unknown’, its value was set to missing values. Third, if levels of one modifier prevent the use of another modifier (for example, gestures involving the head cannot be coded for laterality) – in those cases, the modifier was set to ‘not valid’. Fourth, modifiers were removed within gesture actions if there was no variation – so, to include modifiers within gesture actions we required that at least two modifier levels occurred at least five times. Individual gesture cases were removed if they had a missing value in a modifier that would otherwise have had sufficient variation to be considered for analysis within that gesture action. For a summary of modifier levels that showed sufficient cases and variation within gesture actions, see Supplementary Material.

### Resulting Dataset

After preprocessing, we were left with a dataset of 7,752 gesture tokens across 42 gesture actions that occurred at least 10 times (range: 10-871 cases, median: 68 cases). The distribution across field sites can be found in Table 1. ‘Body part signaller’ and ‘Body part contact’ were coded for all 42 gesture actions (even if the latter was regularly ‘None’ or ‘Not Valid’ for some gesture actions); ‘Laterality’ was coded for 27 gesture actions; and ‘Repetition’ was coded for 39 gesture actions.

### Morphs

Gesture actions differ in the degree of modifier variation they can show. For example, given the modifiers we selected from within those available in the coded data, the gesture action ‘Somersault’ can only be done in one specific way. For this and other simple gesture actions with one or two modifiers coded, we could just assign a morph to every combination of levels of the two modifiers. However, other gesture actions, such as ‘Touch’ or ‘Hit Other’, have over 30 different combinations of modifier levels available. In most studies (including our own) this generates too few cases across too many categories, and it is therefore impractical for further analyses to assume that each of these distinct combinations is its own distinct signal.

In total, we had 527 distinct gesture action/modifier combinations in the dataset, which would leave us with 13 morphs of each gesture action on average. Using all possible gesture action/modifier combinations as morphs would also ignore the non-independence of modifier levels – for example, it is conceivable that repeated actions are more likely to occur when using both hands rather than one. We need an approach that can find statistically meaningful combinations of modifier levels within gesture actions that occur at least a certain number of times to rule out noise. We chose to set a threshold of five occurrences of modifier levels and morphs throughout to consider them for further analyses. Higher thresholds would reduce the number of morphs by reducing the considered number of dimensions along which clusters are detected and filtering morphs out subsequently, while a lower threshold would allow for more complex splitting choices but potentially create non-replicable morphs based on rare cases. While this gesture dataset represents the largest currently available, it remains smaller than those typically used in linguistic analysis, and thus this threshold is low as compared to standards in linguistic corpora (Levshina, 2015). The smaller datasets available in animal communication research in general and the large number of rare gesture actions and modifier levels in this dataset necessitated a generous approach at the risk of sometimes including spurious morphs.

Our goal was to identify the smallest number of gesture action/modifier combinations (morphs) to which we can accurately assign each gesture case. This last feature is important: a morph is only useful if any new case that is added to the dataset can be assigned to exactly one morph – the same way any observed gesture can be categorised as one and only one gesture action. For this purpose, we use Bayesian Latent Class Analysis (LCA), a model-based cluster detection algorithm (Li et al., 2018), as implemented in R (R Development Core Team & R Core Team, 2020) using the ‘BayesLCA’ package (White & Murphy, 2016).

Latent Class Analysis (LCA) is a statistical technique used to identify underlying subgroups from a set of observed categorical variables (Lazarsfeld & Henry, 1968). These variables are usually unordered and discrete. Our use of LCA here can be seen as the factorial equivalent to the increasing use of Uniform Manifold Approximation and Projection (UMAP) and t-Distributed Stochastic Neighbor Embedding (TSNE) as dimension reduction algorithms for continuous data in vocal analyses (Thomas et al., 2022). We are interested in identifying latent classes based on the modifier levels within gesture actions. The main assumption of LCA is that the observed variables are generated by a finite mixture of unobserved groups (Bouveyron et al., 2019). Each class is assumed to be mutually exclusive and exhaustive, meaning that each observation can only belong to one class. LCA is a model-based approach, which means that it uses statistical models to identify the best solution (Bouveyron et al., 2019). We assess model fit using the Bayesian Information Criterion (BIC), which penalises models with more parameters (Weller et al., 2020). The LCA model estimates two parameters for a given number of underlying classes: the size of each class, and the conditional probabilities of each data point to belong to each class. LCA assumes that the observed variables are conditionally independent, meaning that there is no correlation between variables once the latent class membership is known. This assumption is violated in our dataset - modifier levels can be mutually exclusive or correlated (for example, only certain body parts can be used bilaterally). Small sample sizes may lead to unstable or inaccurate class attributions in LCA (Nylund-Gibson & Choi, 2018) – one of the reasons we removed gesture actions with low number of cases. Having simple and well-separating covariates can mitigate the impact of low sample sizes (Wurpts & Geiser, 2014). Despite the violation of the conditional independence assumption and smaller than ideal sample sizes, we consider LCA a useful tool for splitting gesture actions further, but researchers should be aware that some of the found cluster solutions for the less well represented gesture actions can be unstable and may fail to replicate in new data. We assume that, with more data, more morphs would be detectable and some of the less-well represented current morphs would change, either because currently excluded modifier levels would become available or because the LCA would split clusters that are currently lumped. We also lack sufficient cases for each individual/gesture type combination to account for individual differences, potentially another source of pseudoreplication. However, the same problem (larger sample size reveals more distinct elements) applies to any repertoire analysis, and the better represented – and thus more stable morphs – are those that also represent the majority of gestures used by chimpanzees. The number of unique modifier levels that were observed for a gesture action determine the maximum number of clusters that could be detected as they determine the number of binary dimensions along which the cluster detection takes place. Thus, a gesture action for which the coding scheme only allowed two different levels of one modifier will be limited to a maximum of two clusters, while one with variation across all modifiers could be split much more finely given sufficient cases.

Our approach was to take all cases for a given gesture action, remove modifiers that did not show sufficient variation (see *Preprocessing*), and one-hot dummy code modifier levels so that each modifier level was represented as 0 or 1 for each gesture case (Hvitfeldt & Silge, 2022). We based the maximum number of morphs into which a gesture action could be split on the number of unique combinations of modifier levels within the gesture action. We then used LCA to determine the fit for each possible number of clusters between one and that maximum value, ten times per possible number of clusters. This last step increases robustness, as different iterations of the LCA could lead to different results, based on different starting conditions (Li et al., 2018). We extracted the BIC as a measure of model fit of the latent solution. The best cluster solution had to fulfil two conditions: a) all clusters had to be deterministic with regard to a set of modifiers, so that new cases can be unambiguously assigned to one of them; b) none of the resulting clusters included fewer than five cases (Weller et al., 2020). If multiple cluster solutions fulfilled these criteria, we chose the one with the lowest BIC.

For the chosen cluster solution, we extracted the cluster assignment for each gesture case. We subsequently tested whether all clusters within a gesture action could be explained exhaustively by any linear combination of modifier levels, by calculating the probability that each modifier level combination occurred in each cluster and the specificity of this modifier level combination to the cluster. If any modifier level combination had a probability of 1 (meaning it occurred in all cases of the cluster) and a specificity of 1 (meaning it only occurred in this cluster and no other), this modifier level combination became the defining rule for the morph. These assignments can therefore be constructed as a decision tree that gets increasingly complex. For example, the gesture action ‘Object Shake’ is split into four clusters - Object Shakes without repetition (with any limb); with repetition and using the foot; with repetition, using one hand; and with repetition, using both hands simultaneously. If a new case is added, it can be immediately assigned. For gesture actions with clusters that were not exhaustively described by any one modifier level combination, we checked whether several modifier level combinations reached perfect probability and specificity (for example, a cluster that combines the use of hand and fingers as body part). If this did not lead to a reproducible ruleset to describe the morph, we left this morph as ‘unexplained’. This only occurred in this data set once and was due to the LCA creating an ‘all other cases’ category.

### Value of Morphs

While it is possible to split gesture actions into morphs based on modifiers, the question remains whether a maximally split repertoire is valuable to researchers. We propose two criteria that would make a given morph valuable for current gesture research: a) if we can establish that there are clear-cut community differences in the usage of (some) morphs of the same gesture action; and/or b) if morphs of the same gesture action have different meanings and reduce uncertainty about the outcome of the interaction. We use the goal or ‘apparently satisfactory outcome’ of the gesture action as a proxy for meaning (following Cartmill & Byrne, 2010; Hobaiter & Byrne, 2014). All goals and their definitions can be found in the Supplementary Material. After establishing the morphs, we calculated community and goal distributions within gesture actions that have at least two morphs. We established whether a morph had less uncertainty than the gesture action in either domain by calculating the information entropy of communities and goals within each morph of a gesture action (Shannon, 1948) using the ‘infotheo’ package (Meyer, 2022). We then permuted the community/goal membership across morphs within a gesture action, and calculated the resulting entropies, with 1,000 repetitions. This permutation procedure establishes the expected entropy of community/goal distributions if the distribution within gesture actions was random. If the observed entropy of a morph is lower than expected (i.e., smaller than at least 950 of 1,000 permutations, at an alpha level of p = 0.05), we assume that the morph split increased our ability to correctly predict the outcome of a gesture from the morphs, rather than just the gesture action. For this analysis, we removed cases where goals could not be established, which is often the case if recipients do not openly react to a gesture or produce a reaction that is not considered a plausible goal (e.g., an attack). We make the assumption that missing goal assignment is randomly distributed across morphs within gesture actions, an assumption that was only violated in 3 out of 42 gesture actions (based on a Chi-square test for known and unknown goals within gesture actions).

As an additional way to test whether morphs increase the predictability of the community or goal, we implemented a naïve Bayes classifier using the ‘naivebayes’ package in R (Majka, 2019). Naïve Bayes classifiers are well-established and implemented, using vectors of feature values (in our case, the gesture action or morph) to predict the correct outcome (here, the goal or community) using Bayes theorem (Eisenstein, 2019). The dataset was split into ten folds, the classifier was trained on 90% of the data, and the correct classification rate of the remaining 10% was noted (Hvitfeldt & Silge, 2022). We assume that if morphs are a relevant split, they should predict goals better than gesture actions alone, either across the whole dataset or within a subset of gesture actions.

## Results

For 12 gesture actions, only one way of performing them was ever present in the dataset after removing rare occurrences of modifier levels. Of the remaining 30 gesture actions, in a further five cases, the Latent Class Analysis extracted one cluster as the most likely solution. Thus, 17 out of 42 gesture actions were represented as a unified single morph across sites in our dataset (in addition to the 19 gesture actions for which insufficient data were available to establish morphs). The remaining gesture actions were each split into between 2-7 morphs (see Table 2 for their distribution). In all, the dataset yielded 115 morphs. Table 3 contains information on all gesture actions, the number of morphs into which they were split, and the modifiers that were used to make the split. For 99 out of 115 morphs, one clear rule of modifier level combinations could be established that would allow new cases to be sorted into these categories. Of the remaining 16 morphs, 15 were combinations of rare cases; e.g., instead of splitting ‘Biting’ based on all contact points, the rare cases of biting the face, hand, and legs were combined into one morph. After removing rare cases and gesture actions with missing information, 7,560 out of 7,749 gesture cases could be assigned to a single morph.

**Table 2:** Distribution of the number of morphs that the 42 gesture actions were split into.

| Number of Morphs per Gesture Action | Frequency |
| --- | --- |
| 1 | 17 |
| 2 | 6 |
| 3 | 7 |
| 4 | 4 |
| 5 | 1 |
| 6 | 5 |
| 7 | 2 |

**Table 3:**
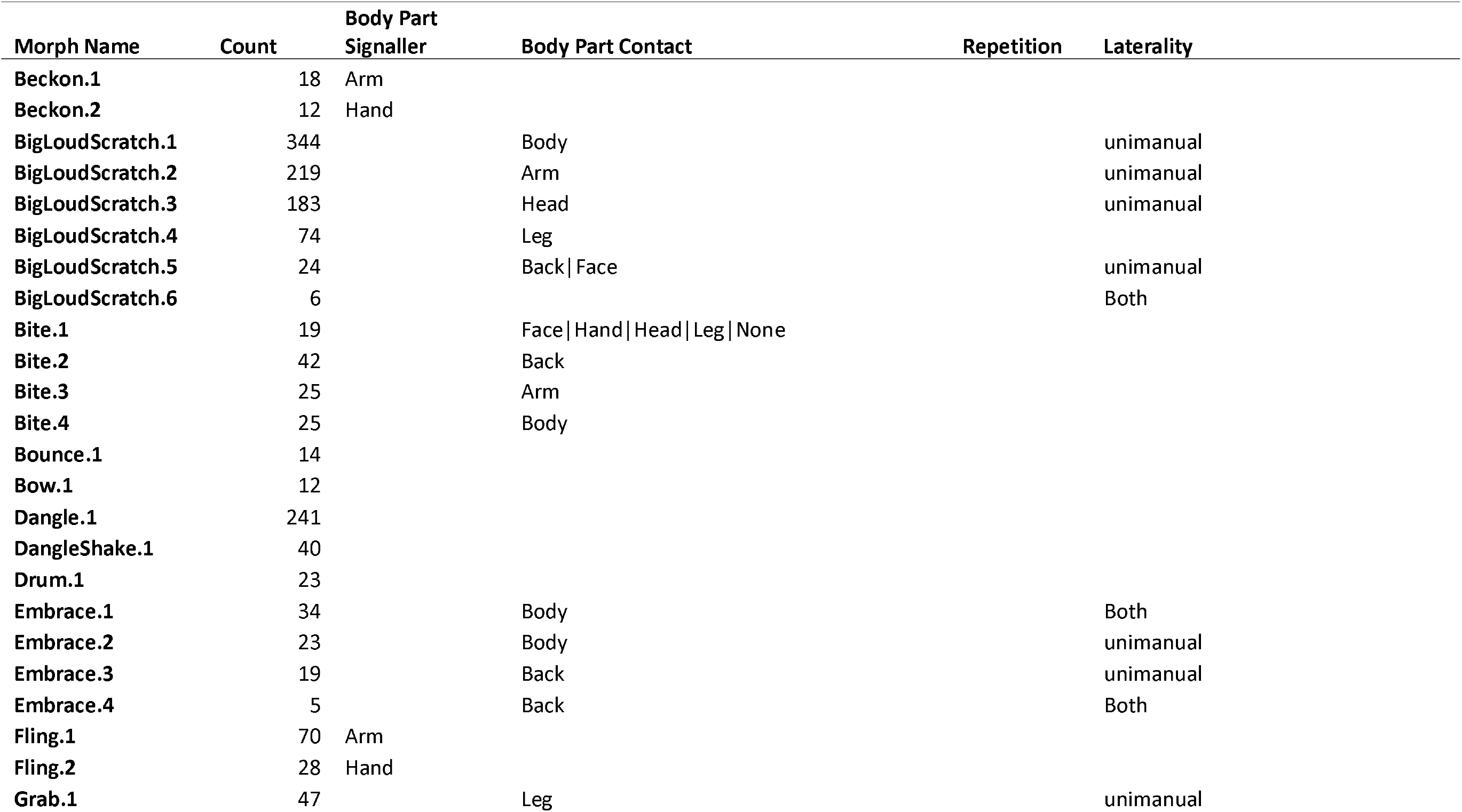
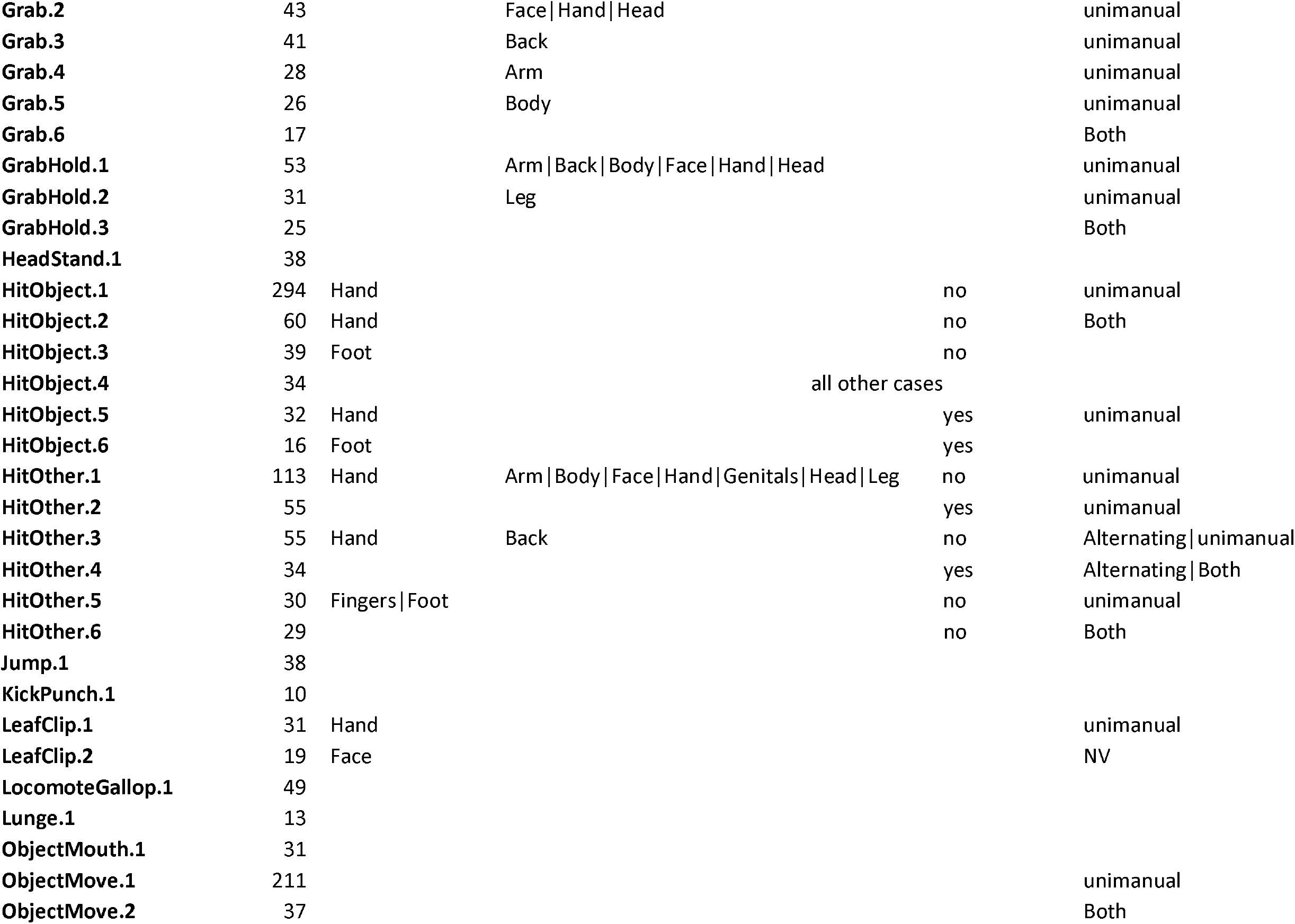
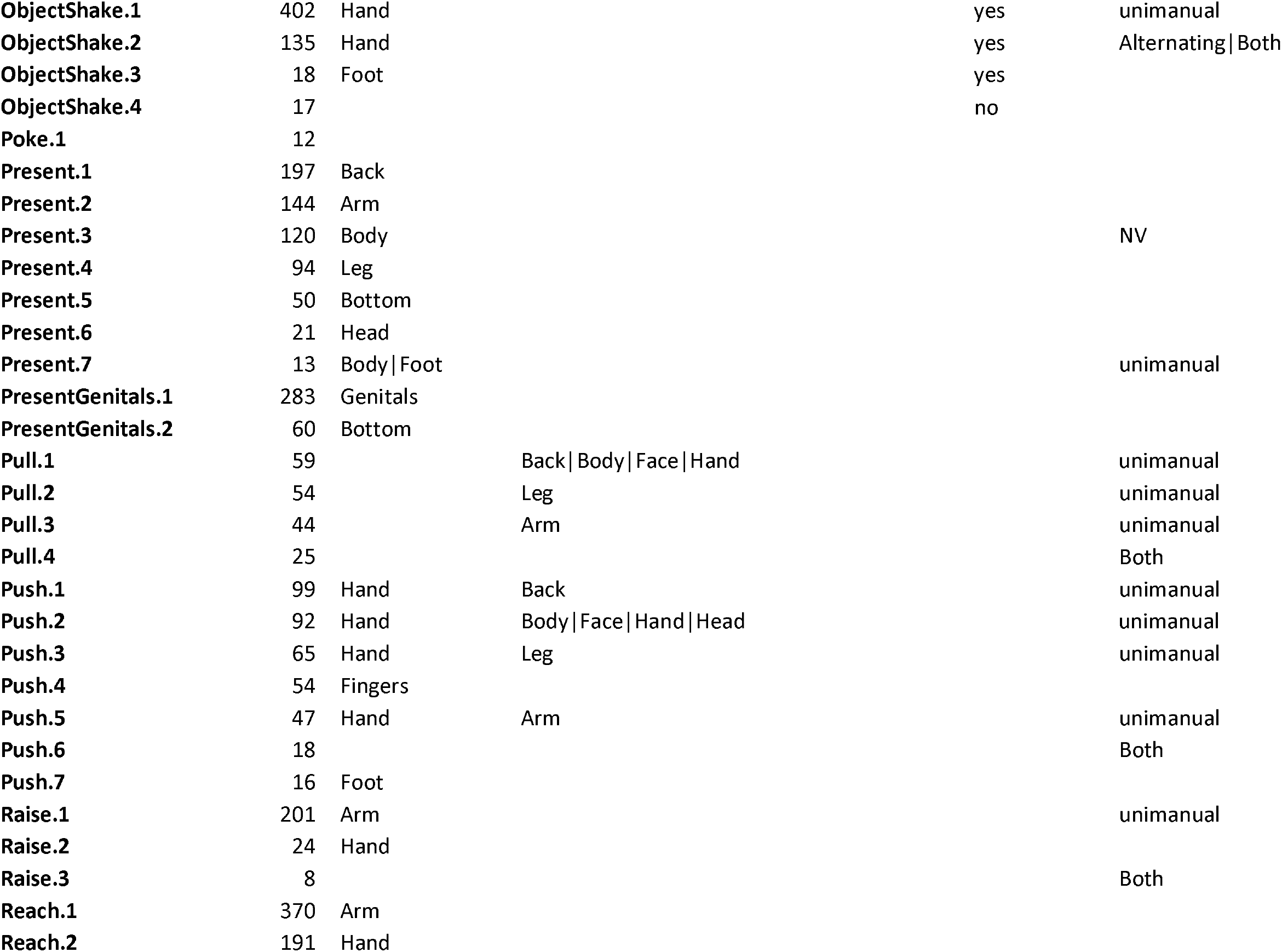
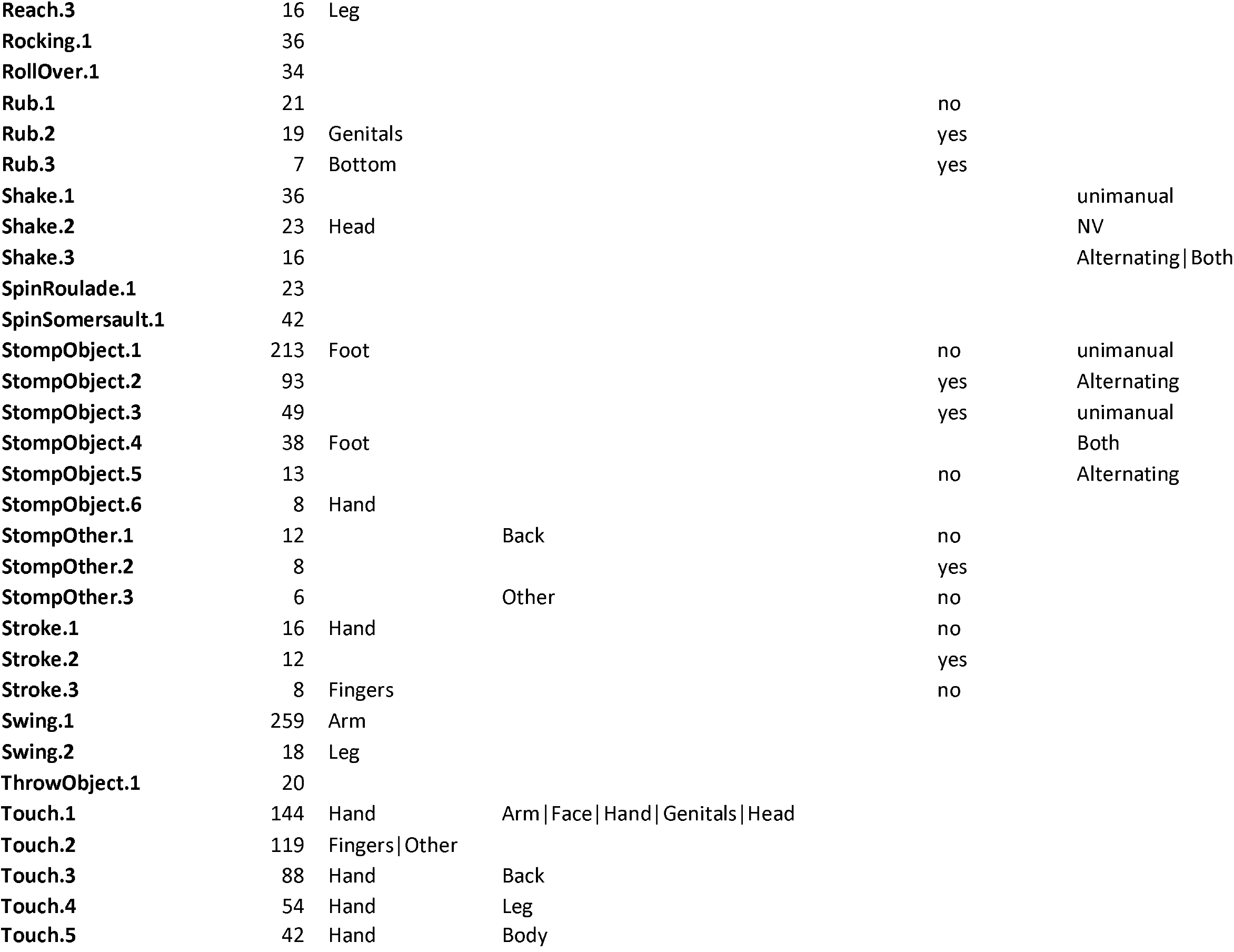
All morphs, and the modifier levels that define them. Dashes indicate that either of those levels would be classed in that morph. NV indicates that the modifier level is not defined within another modifier level (e.g., ‘Laterality’ for gestures using the head)

Splitting of gesture actions was primarily done based on body parts and laterality, as multiple choices were available in many gesture actions (Figure 1). Repetition was less relevant, because most gesture actions occurred either exclusively with repetition or without. Of the 42 gesture actions, 17 (41%) were not split based on any modifier (because there was only a single morph to the gesture action), 9 gesture actions (21%) were categorised as morphs by a single modifier, 11 gesture actions (26%) were split along two modifiers, 4 gesture actions (10%) were split by three modifiers, and 1 gesture action (2%) was split by four modifiers.

**Figure 1:**
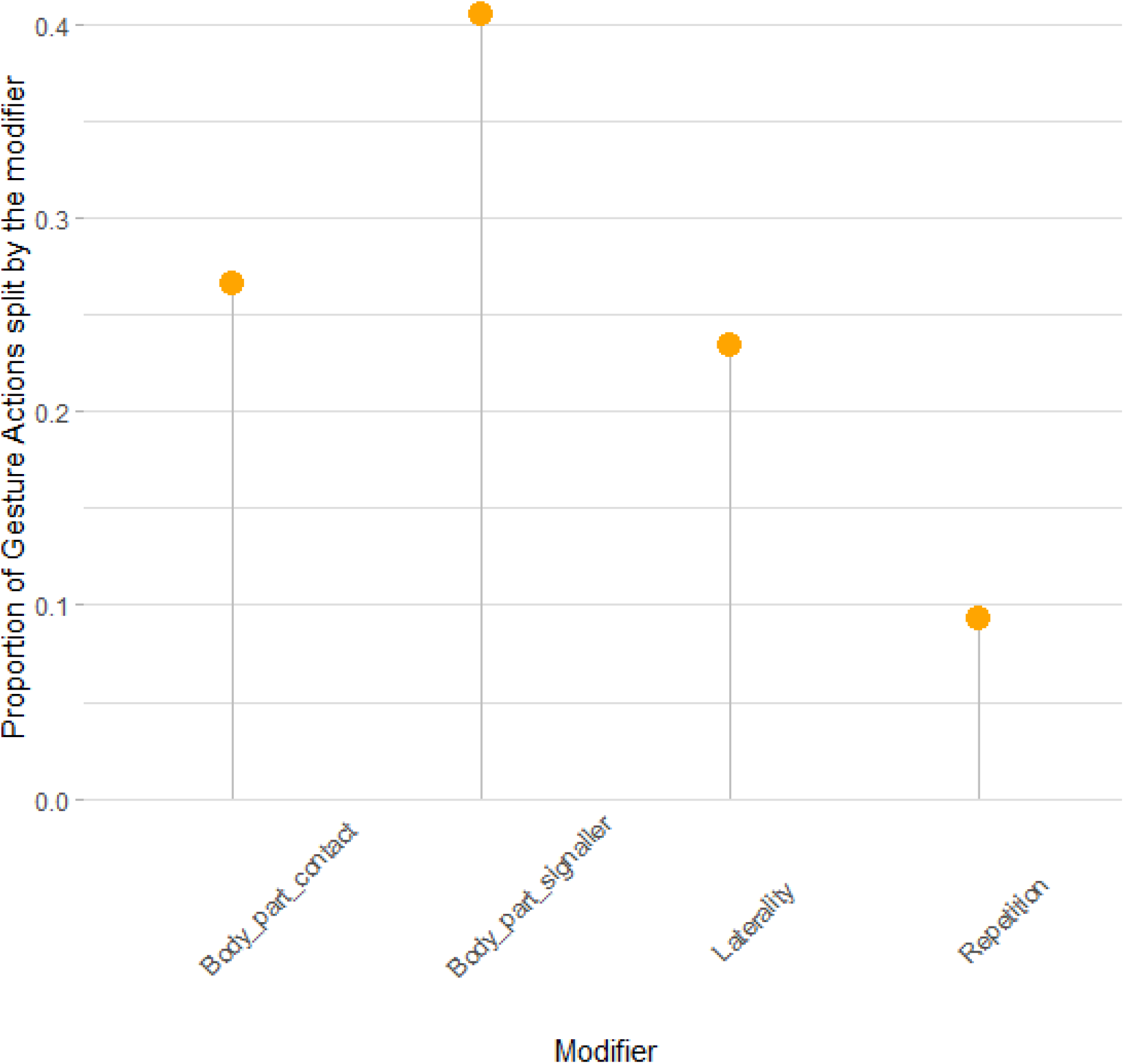
Proportion of Morphs that use each modifier as part of their ruleset.

When looking at the uncertainty of the goal of morphs and gesture actions, we find that in 21 out of 25 gesture actions that were split into morphs, at least one morph existed that had significantly lower entropy (less uncertainty) than would be expected given the distribution of goals in the gesture action itself. This pattern indicates that splitting the gesture action into morphs potentially increases prediction certainty about the meaning. In only 1 out of 21 gesture actions was the morph in question exclusive to play; thus, this decrease in entropy is not driven by the presence of highly predictable play morphs. Throughout the dataset, only 3 out of 115 morphs were exclusive to play (two ‘Stomp Other’ morphs, one ‘Grab’ morph). Similarly, we find that 23 out of 25 gesture actions that were split into morphs had at least one morph that decreased uncertainty about the community in which the morph was observed, indicating the potential for community specific patterns in the representation of morphs in gesture repertoires. Of those, in only two gesture actions were the morphs specific to the Sonso community suggesting that, again, any variation in entropy was not driven by sampling biases.

Using a Naive Bayes classifier, morphs (correct classification = 0.34, expected correct classification at random 0.01) are slightly worse predictors of goals overall than gesture actions (correct classification = 0.37) on the individual gesture level. However, this is not the case across goals (Fig. 3). For 16 out of 25 goals, the morphs actually provide better prediction accuracy, and on the goal-level (treating the average accuracy of each goal as a data point), morphs lead to 32% correct predictions while gesture actions lead to 26% correct predictions. The difference arises because Play, which provides around one third of all data points, is more accurately predicted by the gesture action. The improvement due to using morphs for predictions includes seven goals that would never be predicted correctly if the lumped gesture actions were used as a predictor, likely because they are a secondary goal for a gesture action with one dominant meaning. After splitting the gesture actions into morphs, some of those goals (e.g., Travel, mother-infant communication) are correctly predicted at relatively high rates, indicating that splitting gesture actions more finely does offer important information in those situations. For example, travel is often initiated using Big Loud Scratch gestures (46% of travel initiations); however, the most common goal of that gesture action is to initiate grooming (66% of cases). For some of the morphs, Travel is the most common goal after upsampling, so the classifier correctly assigns this target.

**Figure 3:**
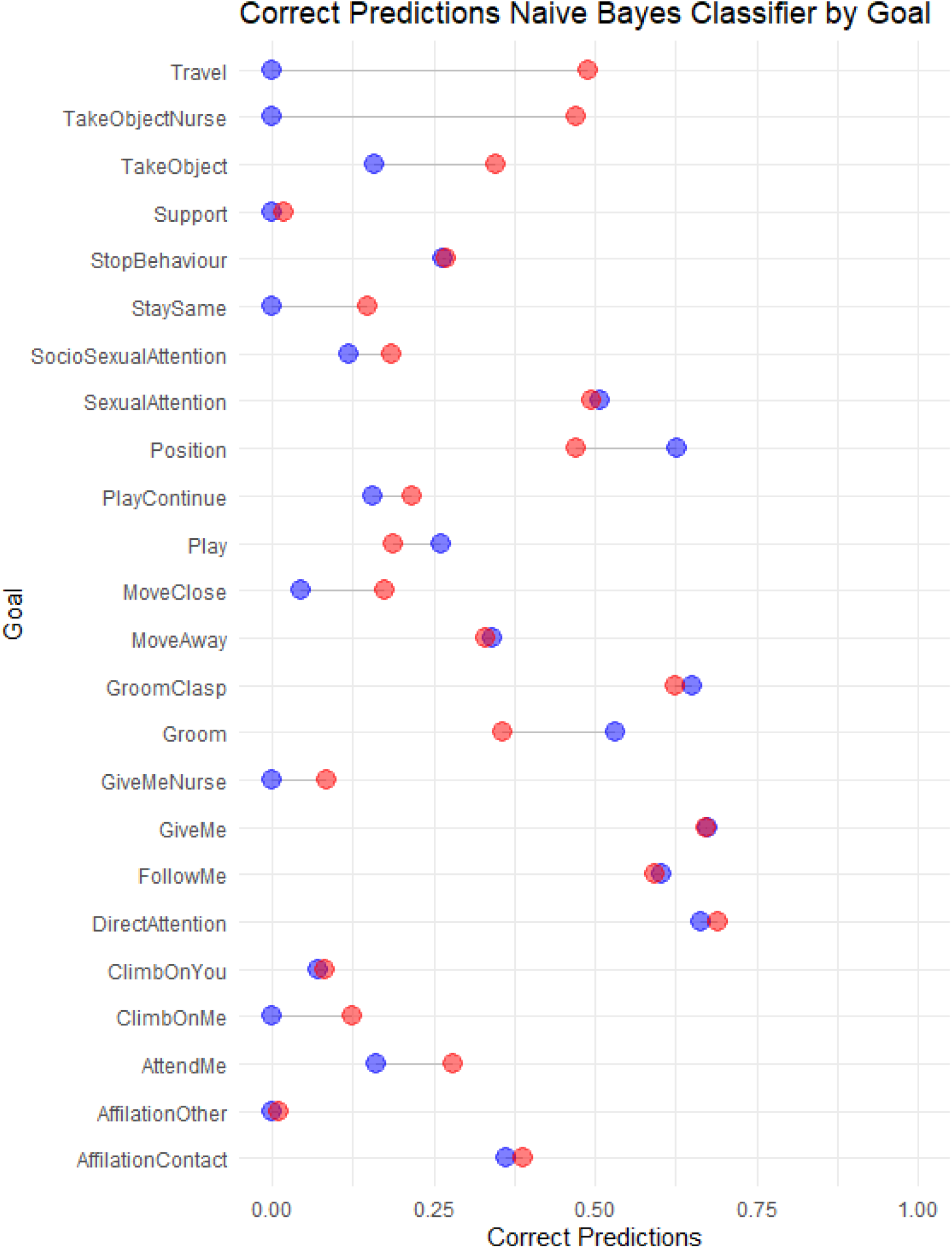
Prediction accuracy in a Naive Bayes classifier for different goals based on gesture actions (blue) and morphs (red). Random prediction accuracy ca. 0.01.

## Discussion

One of the first tasks facing any researcher of animal communication systems is to delineate discrete elements based on their own experience, available data, and perception of the study animal. Historically, there has been a drive to start with a small number of easily discriminated elements (be they facial signals, gestures, or calls), and increasingly split them as researchers become more familiar with variation in species and individual behaviour and get better at recognising nuances that represent variation within, as compared to differences between, signal units, and as new tools become available for analysis. Through this process, signal contexts become clearer. In vocal and facial communication, recent developments in automated and manual feature extraction and classification, respectively, have allowed researchers to extend repertoires of elements or move past them to treat signals as graded and continuous on multiple dimensions (Mielke et al., 2021; Thomas et al., 2022). Here, we show that while gestural repertoires tend to have been considerably larger than those in other channels, chimpanzees may be encoding nuanced information that we are missing using typical levels of lumping in pre-defined gestural repertoires, justifying the process of detailed, bottom-up video coding to understand gestures as a communication system (Grund et al., 2023). We also show that the habit of lumping and splitting elements within a repertoire at different levels of granularity based on non-systematic use of modifiers potentially creates problems for comparative research. Not only does this create imbalances within a particular repertoire that may impact analysis and interpretation – much like treating an assortment of phonemes and words as a single language set – inconsistency across studies often leads to substantial sacrifices of the between-study comparison and replicability often essential to combine data from populations for the species and family-level comparisons used for phylogenetic and evolutionary interpretation (Rodrigues et al., 2021). We argue for consistent and transparent application of lumping and splitting rules along modifying dimensions.

In this study, using an exceptionally large gesture dataset for East African chimpanzees and latent class analysis as a model-based unsupervised classifier, we show that data-based splits exist for most gesture actions that provided sufficient cases, and that the majority of the resulting ‘morphs’ contain information that reduced uncertainty about the goal or chimpanzee community. Splitting here is done purely based on distributions and co-occurrences of specific modifier levels, meaning the approach is consistent across gesture actions, increasing replicability and comparability across studies and species. Thus, rather than splitting ‘Stomp Object’ based on whether it was done with one or two legs and ‘Hit Object’ based on whether it was done once or repeatedly, as was done previously in subsets of this dataset (Byrne et al., 2017; Hobaiter & Byrne, 2011), both are split based on the distribution of the used body part, the repetition, and the laterality (standard modifiers in sign language coding; Kendon, 2004). However, while ‘Stomp Object’ contains alternating use of both limbs with and without repetition, ‘Hit Object’ was not observed alternatingly, leading to a larger number of morphs for the former than the latter. The result is a maximally split dataset within the confines of the observed data and chosen thresholds. The number of morphs we observed (115 from 42 gesture actions with sufficient data) is considerably smaller than the number of all observed combinations of modifiers and gesture actions in the dataset (527), but all morphs occur at least five times and potentially erroneous variation in the modifiers due to coding irregularities was removed by excluding rare occurrences. If they were split, gesture actions fell into 2 to 7 morphs, providing the basis for further investigations of usage, for example in individual or community level variation (c.f., Badihi et al., 2023). The resulting set of gesture morphs allows for more variation and information than represented at the level of the gesture action repertoire, but remains small enough, with sufficient data density per unit, to handle most analyses of gestural communication.

One consistent result of ape gesture research is that – like words in human languages – gestures show means-ends dissociation (Tomasello et al., 1994; Hobaiter & Byrne, 2014): most gesture actions are used for multiple goals, and most goals can be achieved using multiple gesture actions (Graham et al., 2020). Another common result is that there is large overlap across ape species, subspecies, populations, and communities in existing gesture actions and their usage (Byrne et al., 2017; Graham et al., 2018). However, both these findings are based on the assumption that all gesture instances (often termed tokens) that are assigned to one specific gesture action are, in fact, the same communicative signal. Splitting the gesture actions into morphs could potentially reveal differences in meaning and community patterns, enabling us to get a more nuanced picture of primate gesture usage, particularly where a more systematic approach to categorisation allows us to minimise structural coding biases towards the most well-studied species (in primate gesture, typically chimpanzees; Grund et al., 2023; Rodrigues et al., 2021). Here, using morphs rather than gesture actions reduces the uncertainty about the goals and communities associated with some gesture actions: they become more predictable. This is a strong indicator that, while gesture actions see widespread use across meanings and communities, morphs are more specific and potentially a more relevant unit of analysis for some analyses (e.g., Byrne et al., 2017). This leads to improved predictions for many goals when using a Naive Bayes classifier to predict the meaning from the gesture action or morph alone. Crucially, this is not the case across all gesture actions - for around half of them, even though morphs reduce entropy, they do not improve predictability or even add noise when using only one predictor (morph or gesture action, respectively) and a fairly simple classifier that puts high value on the most common observed target to maximise predictions. Disentangling the relationships between morphs or gesture actions and meaning or community membership will be an important task for future research, as will be the question of whether the reduced uncertainty of some morphs is driven by specific modifiers. This finding is equivalent to recent studies showing the depth of chimpanzee cultural diversity by further refining coding schemes for tool-use behaviours (Boesch et al., 2020). It remains a key step forward to recognise that, for many research questions, gesture actions might be insufficiently split to make conclusive statements about variation and flexibility of use.

Importantly, most gesture studies will not have the same level of detail or sample size as this one and may need to lump morphs or even gesture actions (rather than splitting) in order to reach a sufficiently large sample per gesture unit. Even in our dataset, around one third of all coded gesture actions did not occur at least 10 times, the threshold we set for the inclusion of gesture actions in the analysis. What then, can we achieve by splitting the repertoire ever more finely? By showing that modifiers matter and gesture actions can be split more finely, we hope to encourage researchers to code their original data with additional detail and increase the use of replicable and comparable categories and ethograms, which will make the combination and re-analysis of datasets easier (Cartmill & Hobaiter, 2019; Grund et al., 2023; Rodrigues et al., 2021). Further, it is crucial to reiterate that there is no ’correct’ level of granularity in a gesture repertoire. In every communication dataset, there will be some elements that are common enough to split, potentially allowing more nuanced analyses for that subset of data. There will be some research questions that are driven by variation at higher-levels of lumping – for example, looking at the use of different channels of information (tactile, visual, auditory) does not require splitting even by gesture action (e.g., Liebal et al., 2004; Gupta & Sinha, 2019; Dafreville et al., 2021). As long as research methods remain transparent about the level of analysis taking place, fine-grained coded data can be reconstituted into different categories driven by the questions of interest. To an extent, this post-hoc re-classification at a more lumped level already occurs due to data restrictions, with researchers lumping or excluding rare elements where analyses would otherwise become untenable. In the case of dataset driven decisions, we advocate for an approach that creates the ‘most-split’ dataset possible (in the current case, our morphs), and, where lumping is needed, this should occur following clearly defined rules that are established a priori to coding.

An important consideration of our current repertoire of morphs is that they remain – like all repertoires – based on a set of decisions we made at the level of the coding scheme (for example, the types and splitting of modifiers that we included) and at the level of the current analysis (for example, the modifiers included and chosen thresholds). Thus, while substantially *more* bottom-up than other approaches, even more fine-grained levels of splitting are theoretically possible, as would be the inclusion of different modifiers (given a sufficient dataset). Because the latent class analysis is model-based and dependent on initial conditions, where modifier levels are rare, individual morph assignments may change between runs. In practice, however, more fine-grained coding is extremely unlikely across whole gesture repertoires because of the labour-intensive nature of coding and the size of the datasets required. Even with promising new methods to automate the detection and description of ape body-posture within video (e.g., Wiltshire et al., 2023), full automation of gesture coding in video of wild apes remains a substantial challenge. Perhaps more importantly, the continued refinement of gestural repertoires as we continue to study ape gestures and learn more is the scientific process functioning as it should. As our understanding of ape gestures becomes more refined, and as more data become available, it is likely that we will detect more morphs that currently fall below the threshold. This is akin to increasing sample sizes in any description of animal behaviour – but the crucial difference here is that the splitting process can be automatised and repeated following an established set of rules.

There is no correct level of splitting or repertoire granularity; just as phonemes, syllables, and words represent valid levels of linguistic analysis, gesture actions, gesture morphs, and even more finely split units are all valid levels of gesture analysis. But, just as in the study of languages, it is key that a) repertoires are composed of units that are split consistently and by features that are salient (based on recipient responses) to their *users* – here, East African chimpanzees, and b) that researchers tailor their use of different repertoires to their question.

Important claims about the semantic flexibility of great ape gesture and its similarity to human word use (Tomasello et al., 1994; Hobaiter & Byrne, 2014) are based on the assumption that the repertoires applied represent a level of splitting suitable for unit-meaning mapping. Conversely, there remains no evidence for syntactic structures or combinatoriality in ape gesture, despite substantial research effort (Liebal et al., 2004b; Genty & Byrne, 2010; Hobaiter & Byrne, 2011b; Graham et al., 2020), but the detection of structural rules is only possible where signals are parsed into relevant units. In human language, variation in the tone of word production may reflect variation in emphasis or affect (as in English or Arabic) or may fundamentally change word meaning (as in Thai or Cantonese). In gesture, modifiers such as rhythmic repetition, the body part involved in production, or the inclusion and modification of objects may be used to vary emphasis or to change meaning (Kendon, 2004). The systematic description and investigation of modifiers of ape gesture production is an essential step in our understanding of gesture as a communication system. Here, we build on a newly described approach to gesture coding (Grund et al., 2023) that allows us to apply novel analytical approaches to the construction of gesture units (at the level of gesture actions and gesture morphs) derived from the apes’ own gesture usage. We use the largest dataset of East African chimpanzee gesture currently available to define gesture units and repertoires that can be applied to other, smaller, datasets. We show that previously undescribed levels of granularity (gesture morphs) appear to provide additional specificity when exploring gesture meaning and may allow nuanced description of community-level variation in gesture use. In doing so we provide a foundation to develop the study of ape gesture and to redefine comparisons with other communication systems such as human language.

## Data Availability

All scripts and data can be found here: https://github.com/AlexMielke1988/Morph_Repertoire.

## Acknowledgements

AM was funded by a Leverhulme Early Career Fellowship. CH, GB, KEG, CG, and AS were supported by funding from the European Research Council under Gestural Origins Grant No: 802719. KS and CW were supported by funding from the European Research Council under Grant No: ERC_CoG 2016_724608. We thank all the staff of the Budongo Conservation Field Station, its founder Vernon Reynolds, and the Royal Zoological Society of Scotland who provide core funding. We thank the directors of the Kibale Chimpanzee Project for permission to use video data archives. We thank the Uganda Wildlife Authority, the National Forestry Authority, the President’s Office, and the Uganda National Council for Science and Technology for providing research permits and permissions to conduct research in Budongo, Kalinzu, and Kanyawara. The Issa project (GMERC) is grateful for long-term support provided from the UCSD/Salk Center for Academic Research and Training in Anthropogeny (CARTA). We thank the Tanzanian Wildlife Research Institute (TAWIRI), Commission for Science and Technology (COSTECH), and Tanganyika District for permission to conduct research in the Issa Valley.

We thank all the field assistants and local staff across field sites for the decades of work that make this kind of research possible.

