## Supplementary for "Many morphs: parsing gesture signals from the noise"

***Modifiers – Levels, pre-defined lumping decisions, and frequency***

*Body Part Signaller – 16 Levels*

| **Body Part** | **Lumped with** | **Frequency** |
| --- | --- | --- |
| Arm | Shoulder | 1199 |
| Back | - | 197 |
| Body | - | 654 |
| Chest | - | 11 |
| Front | - | 39 |
| Bottom | - | 138 |
| Face | Mouth, Teeth | 216 |
| Fingers | - | 1117 |
| Fist | - | 27 |
| Foot | - | 599 |
| Genitals | Penis, Swelling, Testes, Genital skin | 316 |
| Hand | - | 2876 |
| Hand/Foot | - | 28 |
| Head | - | 48 |
| Knuckles | - | 104 |
| Leg | - | 144 |

*Body Part Contact – 12 Levels*

| **Body Part** | **Lumped with** | **Frequency** |
| --- | --- | --- |
| Arm | Shoulder | 503 |
| Back |  | 586 |
| Body |  | 441 |
| Chest |  | 70 |
| Front |  | 155 |
| Bottom |  | 71 |
| Face | Teeth | 148 |
| Hand | Fingers, Wrist, Knuckles | 138 |
| Leg | Foot | 360 |
| Genitals | Penis, Swelling, Testes, Genital skin | 59 |
| Head | Throat, Neck | 388 |
| None |  | 4778 |

*Repetition of the Minimum Action Unit – 2 Levels*

| **Repetition** | **Frequency** |
| --- | --- |
| No | 5392 |
| Yes | 2243 |

*Laterality – 3 Levels*

| **Laterality** | **Frequency** |
| --- | --- |
| Unimanual (left or right) | 5323 |
| Alternating | 230 |
| Both simultaneously | 499 |

***Gesture actions – Pre-defined lumping decisions and frequencies***

| **Gesture action** | **Lumped with** | **Frequency** | **Modifiers with variation** |
| --- | --- | --- | --- |
| Beckon | - | 32 | Body_part_signaller |
| Big Loud Scratch | - | 871 | Body_part_contact, Body_part_signaller, Laterality |
| Bite | Bite kiss, Bite threat | 163 | Body_part_contact |
| Bounce | - | 14 | - |
| Bow | - | 12 | - |
| Dangle | - | 241 | - |
| Dangle Shake | - | 40 | Repetition |
| Drum | - | 23 | Body_part_signaller, Laterality, Repetition |
| Embrace | - | 88 | Body_part_contact, Body_part_signaller, Laterality |
| Fling | - | 100 | Body_part_signaller |
| Grab | - | 205 | Body_part_contact, Body_part_signaller, Laterality |
| Grab hold | - | 112 | Body_part_contact, Body_part_signaller, Laterality |
| Head stand | - | 38 | Body_part_contact |
| Hit object | Hit Object Object, Hit Object Soft, Hitting Object, Hitting Object Object, Hitting Object Soft | 484 | Body_part_contact, Body_part_signaller, Laterality, Repetition |
| Hit other | Hit Other, Hit Object Other, Hit Other Soft, Hitting Other, Hitting Other Clap, Hitting Other Soft | 322 | Body_part_contact, Body_part_signaller, Laterality, Repetition |
| Jump | - | 38 | - |
| Kick punch | Kick punch at | 10 | Body_part_contact, Body_part_signaller |
| Leaf clip | Leaf Clip Drop | 54 | Body_part_signaller, Laterality, Repetition |
| Locomote gallop | - | 49 | - |
| Lunge | - | 13 | - |
| Object mouth | Object Mouth Attached, Object Mouth Unattached | 31 | - |
| Object move | Object Move Attached, Object Move Unattached, Rake object | 256 | Body_part_signaller, Laterality, Repetition, |
| Object shake | - | 589 | Body_part_signaller, Laterality, Repetition, |
| Poke | Poking | 12 | Body_part_contact, Repetition |
| Present | Present directed | 661 | Body_part_signaller, Laterality |
| Present genitals | Present genitals forwards, Present genitals backwards | 345 | Body_part_contact, Body_part_signaller |
| Pull | Pull directed | 188 | Body_part_contact, Body_part_signaller, Laterality |
| Push | Push directed | 399 | Body_part_contact, Body_part_signaller, Laterality |
| Raise | - | 236 | Body_part_signaller, Laterality |
| Reach | - | 587 | Body_part_signaller, Laterality |
| Rocking | Rocking Bipedal, Rocking Sit | 36 | - |
| Roll over | - | 34 | - |
| Rub | - | 59 | Body_part_contact, Body_part_signaller, Repetition |
| Shake | - | 77 | Body_part_signaller, Laterality, Repetition |
| Spin roulade | - | 23 | - |
| Spin somersault | - | 42 | - |
| Stomp object | Stomping object | 425 | Body_part_signaller, Laterality, Repetition, |
| Stomp other | Stomping other | 31 | Body_part_contact, Body_part_signaller, Laterality, Repetition, |
| Stroke | - | 40 | Body_part_contact, Body_part_signaller, Laterality, Repetition |
| Swing | Swing directed | 279 | Body_part_signaller, Laterality |
| Throw object | - | 20 | - |
| Touch | Touch Long Other | 470 | Body_part_contact, Body_part_signaller, Laterality, Repetition, |

***Goals – Pre-defined lumping decisions and frequencies***

| **Goal** | **Lumped with** | **Frequency** |
| --- | --- | --- |
| Affiliation contact | Affiliation rest contact | 265 |
| Affiliation other | Affiliation distant, Affiliation unclear | 74 |
| Attend me | - | 52 |
| Climb on me | - | 152 |
| Climb on you | - | 62 |
| Direct attention | - | 275 |
| Follow me | - | 525 |
| Give me | Give me active, Give me passive | 379 |
| Give me nurse | - | 38 |
| Groom | Groom you, Groom me | 1120 |
| Groom clasp | - | 91 |
| Move away | - | 263 |
| Move close | - | 87 |
| Play start | - | 2080 |
| Play continue | Play change | 334 |
| Position | - | 286 |
| Sexual attention | Sexual attention me, Sexual attention you | 547 |
| Socio sexual attention | Socio-sexual attention me, Socio-sexual attention you | 132 |
| Stay same | - | 189 |
| Stop behaviour | Stop behaviour here, Stop behaviour stay | 202 |
| Support | Support me, Support you | 22 |
| Take object | Take object passive | 29 |
| Take object nurse | - | 38 |
| Travel | Travel me, Travel you | 66 |
| Other | - | 8 |
| Unknown | - | 433 |

***Interobserver tests***

| **Variable** | **Test** | **Result** |
| --- | --- | --- |
| Gesture action | Cohen’s Kappa | 0.839 |
| Goal | Cohen’s Kappa | 0.786 |
| Repetition count | Cohen’s Kappa | 0.952 |
| Body part signaller | Cohen’s Kappa | 0.856 |
